# Polymorphism of fecundity genes (*BMP15* and *GDF9*) and their association with litter size in Bangladeshi prolific Black Bengal goat

**DOI:** 10.1101/2020.01.08.899443

**Authors:** A. Das, M. Shaha, M. Das Gupta, Avijit Dutta, O. F. Miazi

## Abstract

Identification of prolificacy associated genetic markers remains vital in goat breeding industry since an increase in litter size can generate significant profit. Black Bengal is a prolific goat breed in Bangladesh. There are no inland reports on polymorphisms associated with fertility of Black Bengal goats in Bangladesh. In this study, we investigated two major fecundity genes-bone morphogenetic protein 15 (*BMP15*) and growth differentiation factor 9 (*GDF9*) in order to detect any possible mutations in these two genes in Bangladeshi Black Bengal goats. We identified six single nucleotide polymorphisms (SNP), of which five (C735A, C743A, G754T, C781A, and C808G) in *BMP15* exon 2 and one (T1173A) in *GDF9* exon 2. We also studied their association with litter size. Association analysis results show that polymorphism at the 735, 754 and 781 nucleotide positions of *BMP15* exon 2 had significant association with litter size in Black Bengal goat. The effect of parity was also highly significant (p <0.001) on litter size. This study explored, for the first time, SNP loci in fecundity genes in Bangladeshi prolific Black Bengal goats. Further studies with a high number of genetically unrelated animals for assessing the association of these loci and others in the fecundity genes with litter size may be useful.

## Introduction

Prolificacy (average number of kids born per kidding) is an essential economic trait in goats. This trait abides vital attention to the animal breeders since an increase in offspring size can yield substantial gains in profit. Prolificacy in domestic species genetically influenced by multiple genes called fecundity genes (Gootwine, 2020). *BMP15* and *GDF9* are two candidate fecundity genes among the several detected in goat so far (Ahlawat et al. 2015). Several studies reported the correlation between prolificacy and genetic polymorphisms in *BMP15* and *GDF9* genes in different goat breeds around the globe, including Black Bengal goat in India (Polley et al. 2009; Hadizadeh et al. 2014; Ahlawat et al. 2015; Ahlawat et al. 2016; Heikal and El Naby 2017; Jalbani et al. 2017; Mishra et al. 2017; Wang et al. 2017; Ghoreishi et al. 2019; Wang et al. 2019; Yue et al. 2019).

In Bangladesh, with a census size of 25.7 million goat stands for the third largest livestock species. Most of the goats in Bangladesh are Black Bengal goat (>90%) (Husain et al., 1998), reared mainly by landless, small-scale farmers. Black Bengal goat is a dwarf goat breed and is well-known for its high prolificacy, meat quality and skin quality (Miah et al. 2016). The average litter size of this breed ranges from 1.50 to 2.13 (Choudhury et al. 2012; Paul et al. 2014; Miah et al. 2016). This breed also renders a significant source of red meat (25%) production in Bangladesh (Amin et al. 2000). Considering its importance, increasing the numbers of black Bengal goat has invariably been the central breeding goal for selective goat breeding program in Bangladesh.

The use of molecular markers in animal breeding has added benefits over conventional breeding techniques. The advancement in molecular genetics has led to the detection of DNA markers with considerable effects on the traits of economic importance. To the best of our knowledge, no screening attempted to identify polymorphisms in fecundity genes in goats of Bangladesh, using DNA based technologies. Therefore, this study aimed at the detection of genetic variants in two fecundity genes (*BMP15* and *GDF9*) and their relationship with litter size in the only recognized native goat breed in Bangladesh, Black Bengal.

## Materials and methods

### Experimental animals and sample collection

In the present experimental study, a total 40 Black Bengal does with twinning record in first three parities were chosen from Hathazari Government Goat Farm, Chattogram district of Bangladesh. Selected Black Bengal does kidded in 2016–2017, there are data on litter size at the first, second and third parity. There were no selection on litter size or other fertility traits in the farm over previous years. The average age and litter size of the does were 38.9 months and 2.48, respectively. These animals were maintained under the standard conditions of management, health, and nutrition.by the farm management.

Blood samples (approximately 10 ml) were collected aseptically from the jugular vein in a vacutainer tube containing ethylene diamine tetraacetic acid (EDTA) as the anticoagulant. All samples were delivered to the Poultry Research and Training Center (PRTC) laboratory at Chattogram Veterinary and Animal Sciences University using an icebox.

### DNA extraction, amplification and DNA sequencing

Genomic DNA was extracted from the blood samples using GeneJET Genomic DNA Purification kit (Thermo Scientific, Waltham, Massachusetts, USA) according to the manufacturer’s instructions. The extracted DNA samples were stored at -20°C prior to being used in Polymerase chain reaction (PCR).

PCR was performed to amplify segments of *BMP15* exon 2 and *GDF9* exon 1 and exon 2. The sequences of primers and amplicon sizes of PCR products are presented in Table1. PCR amplifications were performed in a final reaction volume of 25 μL on a Thermo-cycler (2720 Thermal cycler, Applied Biosystems, USA). The PCR reaction was run under the following thermal condition: initial denaturation for 1 min at 94 °C; 30 cycles of denaturation at 94 °C for 45 s, annealing at 60°C for 45 s, extension step at 72 °C for 45 s with a final extension at 72 °C for 5 min. The amplified PCR product was electrophoresed by running 10 µl of through a 2% agarose gel stained with ethidium bromide (0.5µg/ml) (Sigma Aldrich, USA). The specific sizes of the fragments were distinguished by using 1kb plus DNA ladder (O,GeneRuler, 1 kb Plus, Thermo Scientific Fermentas) in the gel and visualized by a UV transilluminator gel-documentation system (BDA digital, Biometra GmbH, Germany).

In order to detect any possible mutations in exons of *BMP15* and *GDF9* genes, amplified PCR products for 40 samples (one for each animal) were bidirectionally sequenced using BigDye Terminator v. 3.1 (ThermoFisher Scientific, Waltham, MA USA) cycle sequencing protocol by Macrogen Co., Korea.

### Sequence analysis

Nucleotide sequence data were edited and analysed by MEGA version 10.0.5 (Kumar et al. 2018). The detected polymorphisms (SNPs) were compared to the *Capra hircus* nucleotide database in NCBI Genbank using BLAST (Altschul et al. 1990).We used Polyphen-2 (Adzhubei et al. 2013) to predict the possible impact of mutations (based on amino acid substitution) on the structure and function of the respective proteins.

### Statistical analysis

We used SHEsis online platform (http://analysis.bio-x.cn) (Shi and He, 2005) to calculate genotype frequencies, allele frequencies and χ2 values for Hardy–Weinberg equilibrium (HWE) test. The deviation from HWE for each polymorphism was tested using the Hardy– Weinberg law (Gorlov et al. 2018; Yue et al. 2019). Heterozygosity (He), polymorphism information content (PIC) and effective allele number (Ne) for each polymorphism were estimated employing an online computing software (http://www.msrcall.com/Gdicall.aspx). For PIC, following classifications was used i) low polymorphism if PIC value <0.25, ii) moderate polymorphism if PIC value ≥ 0.25 to ≤0.50 and iii) high polymorphism if PIC 0.50.

A linear mixed model was employed to analyse the association of polymorphisms in BMP15 and GDF9 gene with litter size using SPSS 25 statistical software, SPSS Inc., Chicago, IL, USA. The mixed model included fixed effects genotypes and parity (1, 2 and 3). Litter size data from consecutive parities in the same individual were considered as repeated measures.

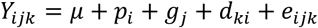

where, *Y*_*ijkl*_ is the phenotypic value of litter size, *μ* is the overall population mean, *p*_*i*_ is the effect of parity, *g*_*j*_ is the effect of genotype, *d*_*ki*_ is a normally distributed random variable with mean zero and variance *σ*^2^_*d*_ corresponding to doe *k* in parity *i* and *e*_*ijk*_ is the random error. The interactions between fixed effects were not significant hence excluded from the model. Mean separation procedures were performed by using the least significant difference test.

## Results

### Sequence results and SNP detection

The sequences were submitted to NCBI Genbank with the accession number of MN401415 and MN401414 for *BMP15* and *GDF9*, respectively. In a comparison of coding sequence sequences of caprine *BMP15* and *GDF9* genes that were recorded in NCBI GenBank (*BMP15*, NM_001285588.1; *GDF9*, NM_001285708.1), we identified five SNPs (G735A, C743A, G754T, C781A, and C808G) in the *BMP15* exon 2 and one SNP (T1173A) in *GDF9* exon 2 (Fig.1). Five of these polymorphisms leading to amino acid substitutions (Table 2). The predicted possible effect of identified polymorphisms on the structure and function of the functional protein revealed G754T and T1173A SNP loci have a significant impact on the coding BMP15 and GDF9 peptide, respectively (Fig.2). The C781CA mutation deduced to cause a moderate change while remaining two mutations did not show significant effects on the functional BMP15 protein.

**Table 1.**
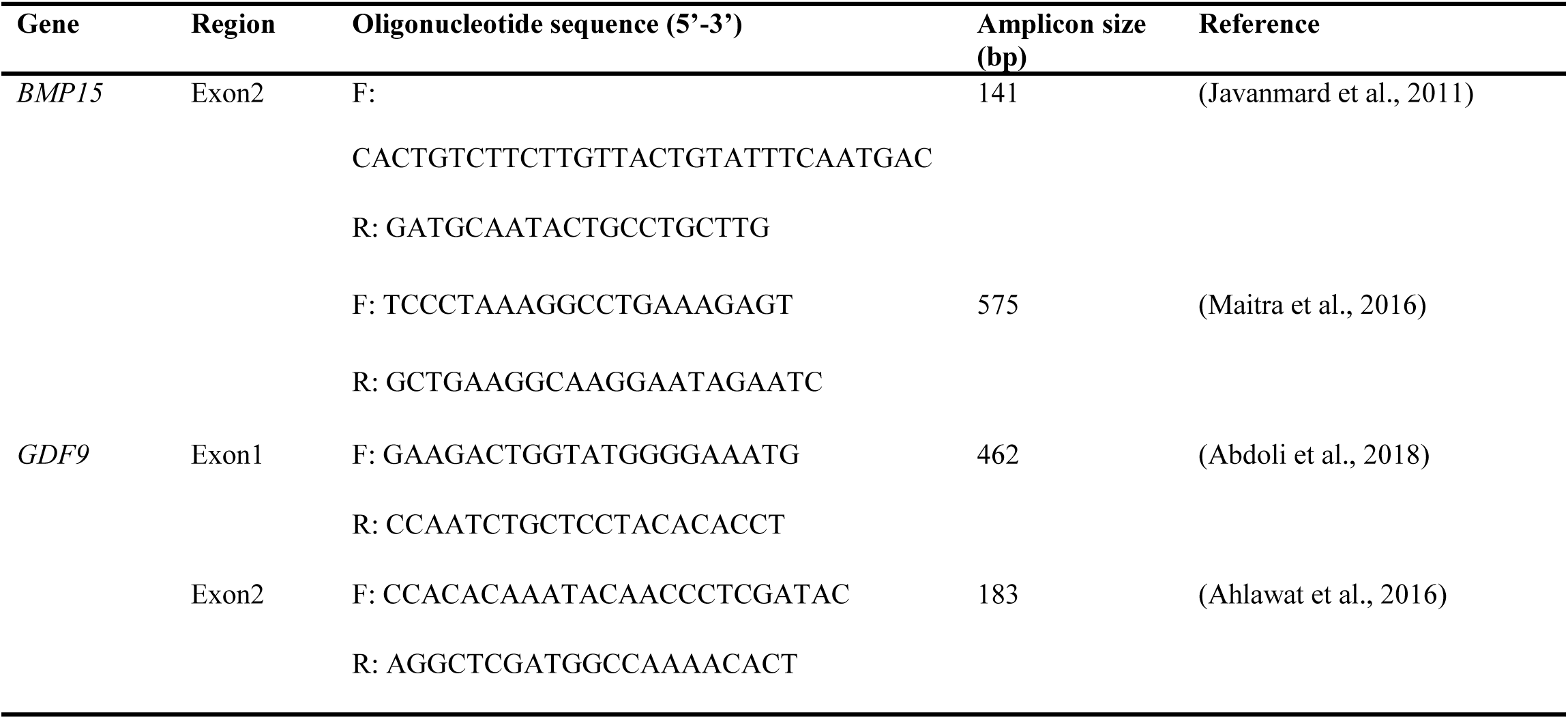
List of primer used to amplify specific segment of *BMP15* and *GDF9* gene.

**Table 2.**
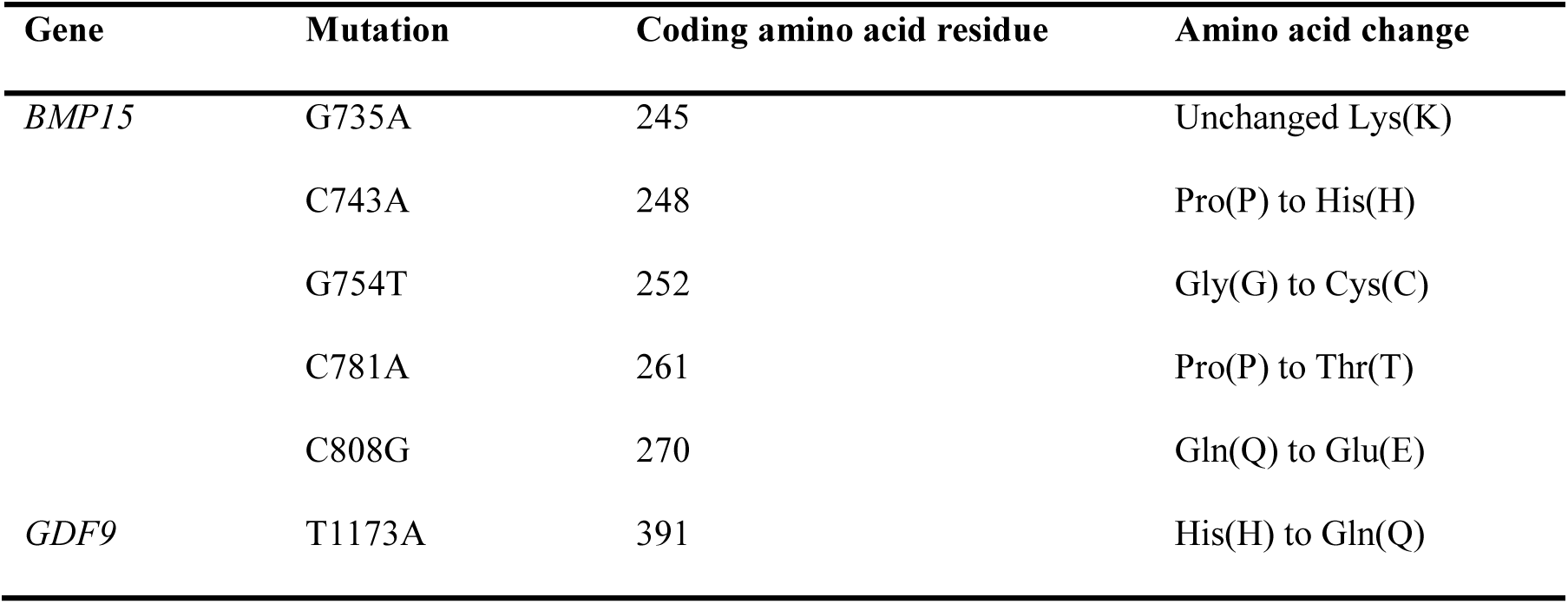
The identified mutations in *BMP15* and *GDF9* within the Bangladeshi Black Bengal Goat breed

**Fig.1.**
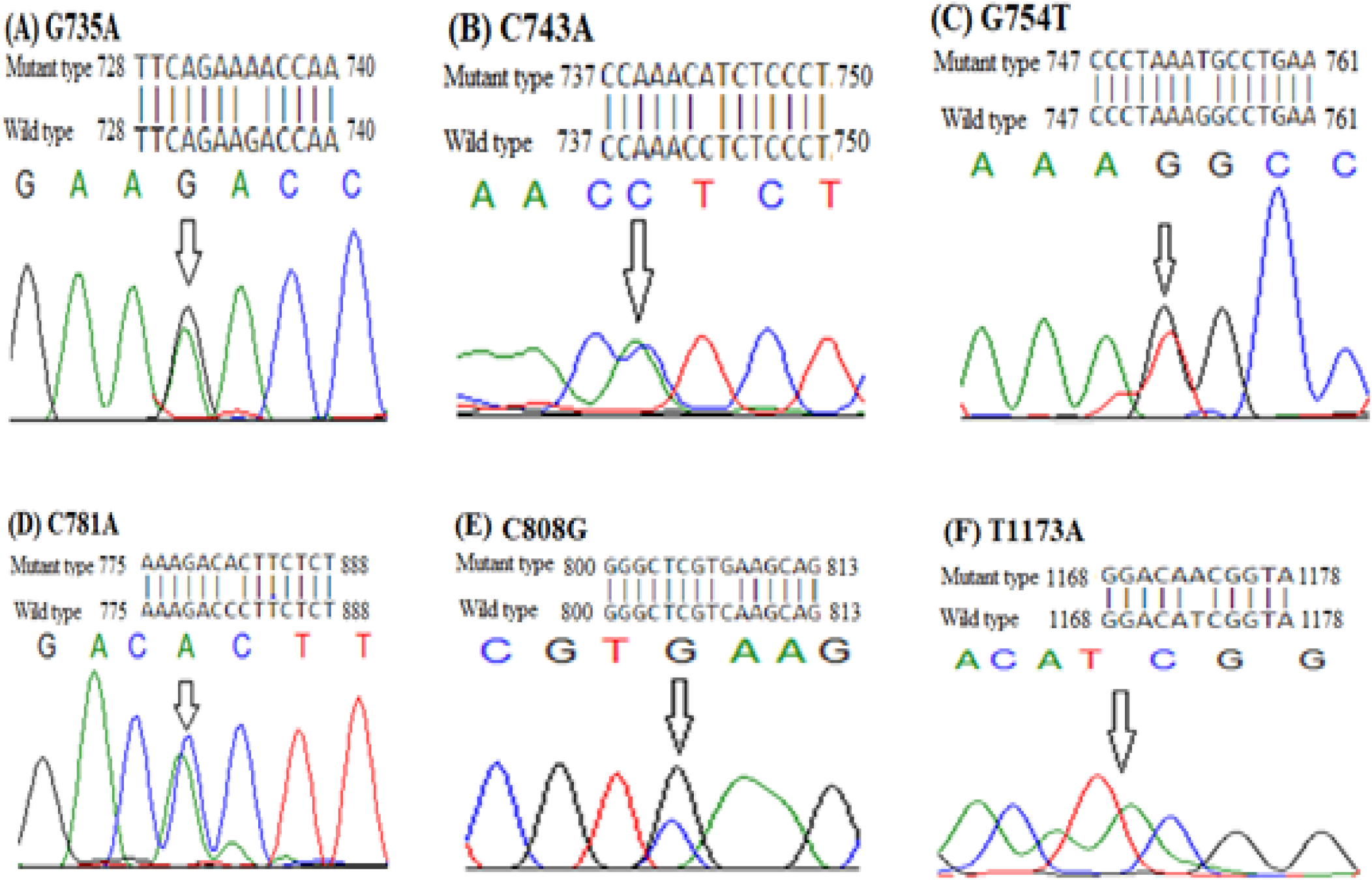
Sequencing chromatograms of the detected SNPs in *BMP15* and *GDF9* genes in Black Bengal goat. **A, B, C, D, and E** represent the SNP loci G735A, G743A, G754T, C781A, and C808G in exon 2 of the *BMP15* gene, respectively. **F**, represents the SNP T1173A **in** exon 2 of the *GDF9* gene. Positions of the mutations are based on the full sequences of the BMP15 and GDF9 gene (Gene IDs: <u>1008612</u>33 <u>and 1008612</u>33, respectively).

**Fig.2.**
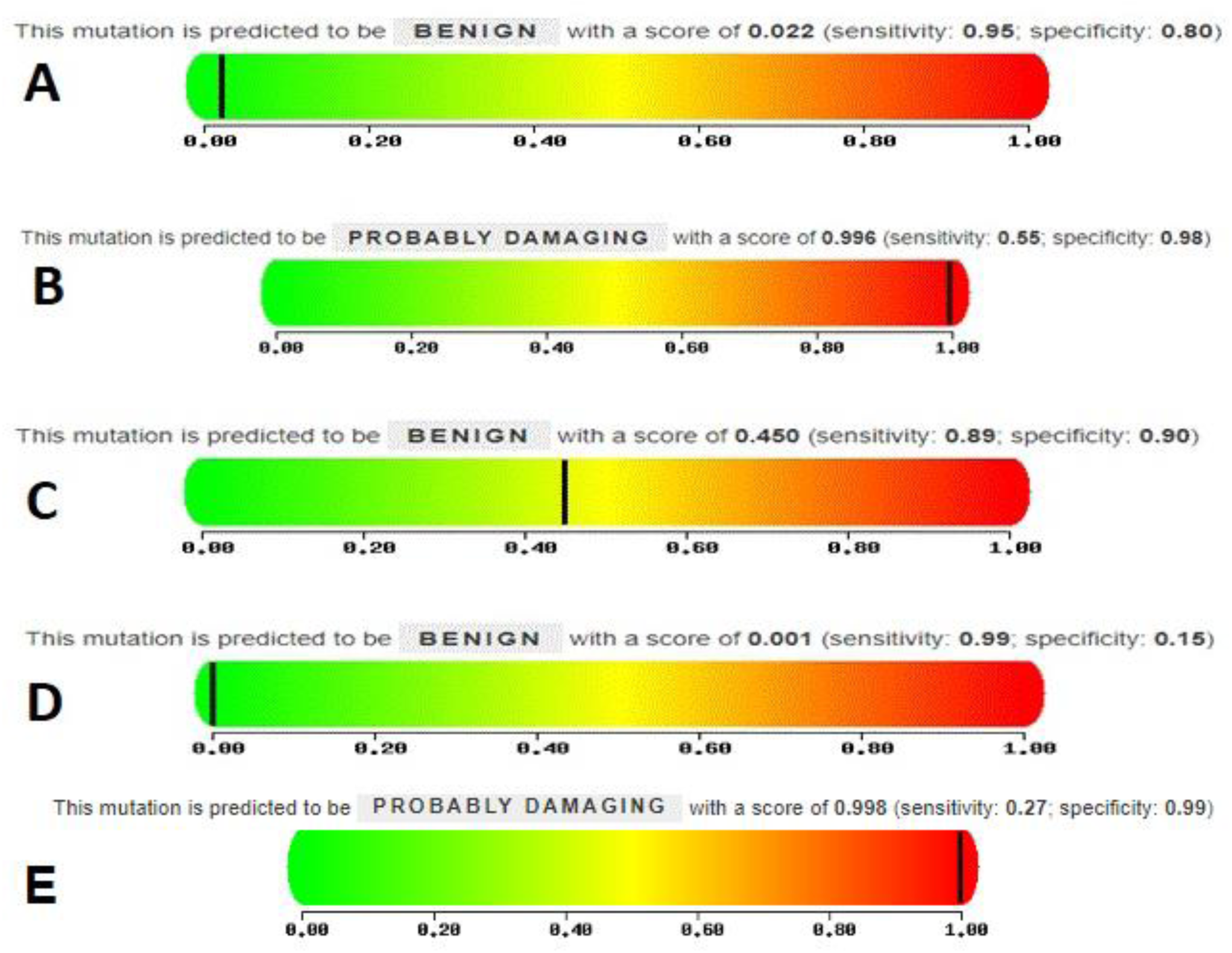
Predicted effects (using Ployphen2) of five SNP loci. **A, B, C**, and **D** represent the effect of C743A, G754T, C781A, and C808G on the functional BMP15 protein, respectively. **E**, represents the effect of T1173A on the functional GDF9 protein.

### Genetic parameters for the detected SNP loci

The genotypic and allelic frequencies, PIC, HE, Ne and χ2 values for detected polymorphisms in two fecundity genes are presented in Table 2. The G735A of *BMP15* showed all the three possible genotypic combinations while remaining SNPs recorded with two genotypes. All tested goats expressed heterozygous genotype at C808G of *BMP15* gene. Genotype frequencies for most of the detected polymorphisms were significantly different from the expectations of Hardy-Weinberg equilibrium in Black goat population in this study (P < 0.01), except G735A and G754T in *BMP15* gene. Three polymorphisms in *BMP15* genes (C743A, C781A and C808G) and one in *GDF9* gene (T1173A) were found to moderate polymorphic according to the classification of PIC. The remainder of the detected polymorphisms was found to be low polymorphic.

### Association between SNP loci and litter size

The least squares means and standard error for litter size of different genotypes at the various loci of BMP15 and GDF9 genes are presented in Table 4. The effect of genotype was significant only for locus G735A (p <0.01), G754T (p <0.05) and C781A (p <0.001) in *BMP15* gene. Parity had a highly significant effect on litter size in tested Black Bengal goat (p <0.001). However, the interaction effect of parity and genotype had no significant effect on litter size. We were unable to test the C808G SNP in *BMP15* gene for association analysis since that had only one genotype in the study population.

**Table 3.** Genotype and allele frequencies of six SNP loci in two fecundity genes in Bangladeshi B 322 lack Bengal Goat (N=40).

| Gene | Loci | Genotype frequency | Allele frequency | PIC | He | Ne | H-W test ($\chi^2$) | p value |
| --- | --- | --- | --- | --- | --- | --- | --- | --- |
| *BMP15* | G735A | GG-0.625 | G-0.762 | 0.237 | 0.361 | 1.567 | 2.317 | 0.127 |
|  |  | GA-0.275 | A-0.237 |  |  |  |  |  |
|  |  | AA-0.100 |  |  |  |  |  |  |
|  | C743A | CC-0.300 | C-0.650 | 0.351 | 0.455 | 1.834 | 11.597 | 0.000 |
|  |  | CA-0.700 | A-0.350 |  |  |  |  |  |
|  |  | AA-0.0 |  |  |  |  |  |  |
|  | G754T | GG-0.700 | G-0.85 | 0.222 | 0.255 | 1.342 | 1.245 | 0.264 |
|  |  | GT-0.300 | T-0.15 |  |  |  |  |  |
|  |  | TT-0.0 |  |  |  |  |  |  |
|  | C781A | CC-0.450 | C-0.725 | 0.319 | 0.398 | 1.663 | 5.755 | 0.016 |
|  |  | CA-0.550 | A-0.275 |  |  |  |  |  |
|  |  | AA-0.0 |  |  |  |  |  |  |
|  | C808G | CC-0.0 | C-0.500 | 0.375 | 0.500 | 2.000 | 40.000 | 0.000 |
|  |  | CG-1.000 | G-0.500 |  |  |  |  |  |
|  |  | GG-0.0 |  |  |  |  |  |  |
| *GDF9* | T1173A | TT-0.300 | T-0.300 | 0.331 | 0.420 | 1.724 | 39.999 | 0.000 |
|  |  | TA-0.700 | A-0.700 |  |  |  |  |  |
|  |  | AA-0.0 |  |  |  |  |  |  |

*PIC*, polymorphism information content; *He*, effective number of heterozygosities; *Ne*, effective number of alleles.

**Table 4.** Parity wise least squares of litter size for different genotypes at different lo 325 ci of *BMP15* and *GDF9* genes.

| Gene | Loci | Genotype | N | Litter size (Mean±SE) |  |  | Effect of genotype |
| --- | --- | --- | --- | --- | --- | --- | --- |
|  |  |  |  | Parity 1 | Parity 2 | Parity 3 |  |
| *BMP15* | G735A | GG | 25 | 1.92±0.11 | 2.64±0.11 | 2.88±0.11 | ** |
|  |  | GA | 11 | 1.82±0.16 | 2.36±0.16 | 2.55±0.16 |  |
|  |  | AA | 4 | 1.75±0.27 | 2.00±0.27 | 2.25±0.27 |  |
|  | C743A | CC | 12 | 2.00±0.16 | 2.58±0.16 | 2.75±0.16 | NS |
|  |  | CA | 28 | 1.82±0.11 | 2.46±0.11 | 2.71±0.11 |  |
|  | G754T | GG | 28 | 1.82±0.10 | 2.39±0.10 | 2.64±0.10 | * |
|  |  | GT | 12 | 2.00±0.16 | 2.75±0.16 | 2.91±0.16 |  |
|  | C781A | CC | 18 | 1.67±0.12 | 2.39±0.12 | 2.39±0.12 | *** |
|  |  | CA | 22 | 2.04±0.11 | 2.59±0.11 | 3.00±0.11 |  |
| *GDF9* | T1173A | TT | 12 | 1.75±0.16 | 2.42±0.16 | 2.75±0.16 | NS |
|  |  | TA | 28 | 1.89±0.10 | 2.57±0.10 | 2.71±0.10 |  |

N, number of observations; \*, P<0.05; \*\*, P<0.01; \*\*\*, P<0.001; NS, Not significant.

## Discussion

We herein reported nine polymorphic sites in two well-studied fecundity genes namely *BMP15* and *GDF9* in Black Bengal goat, the only prolific goat breed in Bangladesh. The genetic parameters of the detected loci and their association with the litter size in the Black Bengal goat were also analysed. The G735A locus of *BMP15* gene recorded with three possible combinations of genotypes. However, for the remaining five SNP loci, we did not observed any homozygous mutant genotype in the study population. One plausible reason is that these mutations are inactivating mutations in homozygote form, therefore does carrying two copies of mutated alleles are sterile and are expected to remove from the reproductive flock. Till date, reports on infertility among Black Bengal goat are scarce. Therefore, absence of homozygote mutant does for these five loci in the Black Bengal goat, similar to the situation that was observed for Small-Tail Han sheep, Tunisian-Barbarine sheep and Markhoz goat (Chu et al. 2007; Lassoued et al. 2017; Ghoreishi et al. 2019).

In this study, three SNPs in *BMP15* gene viz. C743A, G754T and C781A were found to be novel based on an extensive literature search. Remaining two SNPs identified by us in *BMP15* gene viz. G735A and C808G have also been reported in Black Bengal goats in the neighbouring country India (Ahlawat et al. 2013; Ahlawat et al. 2015; Maitra et al. 2016). The SNP in *GDF9* gene (T1173A) detected by us also not reported so far in goat. However, several studies reported polymorphisms in the flanking sequences of *GDF9* gene (Ahlawat et al. 2015; Shokrollahi and Morammazi 2018; Yue et al. 2019). Polley et al. (2009) who screened Black Bengal goats of India for the presence of mutations in *BMP15* and *GDF9* genes already associated with prolificacy in sheep and reported the wild type genotype only. Similar findings have also been reported in five Indian goat breeds in addition to Black Bengal (Ahlawat et al. 2013). These findings differ with the findings of the present study since we screened for goat specific markers. The five non-synonymous (C743A, G754T, C781A, C808G and T1173A) goat specific mutations we detected in the present study might despite the necessity of tested in larger population be valuable for future studies on the molecular and functional effect of mutations on differential prolificacy in Bangladeshi goat breeds. The molecular and functional effect of a mutation can be neutral, harmful or beneficial depending on their context or location (Williams, 2016). In our study, two of the identified polymorphisms (G754T and T1173A) predicted to have a significant effect on resulting protein sequences.

Results of association study showed genotypes at three loci in *BMP15* gene had a significant effect on litter size in tested Black Bengal goat (p= <0.05). Significant effect of genotypes on litter size have been reported in Liaoning cashmere goats and Markhoz goat (Ghoreishi et al. 2019; Yue et al. 2019). In contrast, Ahlawat et al. (2015) and Ahlawat et al. (2016) reported a non-significant effect of genotypes on litter size in Indian Black Bengal goat. Heterogeneity among studies investigating effects on mutations on prolificacy in sheep has also been shown by Gootwine (2020). Presence of more than one segregating mutation in a major gene (*BMP15*) for high prolificacy in the present study is in line with those observed in some prolific sheep breed. For example, both FecXB and FecXG mutations in *BMP15* segregate in Beclatare sheep in Ireland (Hanrahan et al. 2004). Similarly, Tunisian-Barbarine sheep carries FecXBar and B5 mutation in *BMP15* gene (Vacca et al. 2010; Lassoued et al. 2017). None of the loci in *GDF9* gene we detected significantly associated with litter size in Black Bengal goat. Gootwine (2020) also showed that estimates of the effects of major genes vary among studies. Ahlawat et al. (2015) reported a highly significant (p<0.01) effect of parity on litter size in Black Bengal goat of India. In the present study, we also observed a highly significant (p<0.001) effect of parity on litter size. Faruque et al. (2010) reported a significantly (p<0.001) lower litter size in first parity than in the seventh parity in Black Bengal goat. Halder et al. (2014) also confirmed that larger litter size In Black Bengal goat is highly influenced by higher parity and higher litter size in previous parity.

Variations of a quantitative trait are controlled by several genes, genetic variants and their interactions. Hence, the detection of polymorphisms that are underlying the differences in a quantitative trait, for instance, litter size remains as a challenge in modern genetics. However, in some prolific sheep breeds, prolificacy inherited as a qualitative trait, rather than quantitative due to presence of major genes with large effects on litter size (Gootwine, 2020). *BMP15, GDF9* and *BMPR1B* are three well-documented candidate genes for litter size in sheep. Till date, no association of *BMPR1B* gene with litter size in goat has been established. However, researchers have explored the association between litter size in goat and polymorphisms in *BMP15* and *GDF9* genes. These two genes are part of the ovary-derived transforming growth factor β (TGF β) that have an integral role as growth factors and receptors in follicular development in the ovaries (Davis, 2005). Both bone morphogenetic proteins (BMPs) and growth differentiation factors (GDFs) have critical role in follicle growth and cell-survival signalling hence causal mechanism underlying the high prolificacy or fertility in female animal (Otsuka et al., 2011). Considering the biological importance of *BMP15* and *GDF9* genes, this study investigated polymorphisms in these genes and their association with the litter size in Bangladeshi prolific Black Bengal goat.

The frequency distribution of genotype in a population is the simplest way to describe Mendelian variation (Mather and Jinks 1977). Other factors such as such matting pattern, random genetic drift, individual survival, reproductive success, and migration also generate genetic variation in a population (Chen et al. 2019). In the current study we sequenced only 40 goats, it would have an effect on the genotypic and allelic frequencies we observed; hence, large population-scale research is required to confirm the genotypic and allelic frequencies in Black Bengal goat of Bangladesh.

## Conclusion

This is the first study exploring polymorphisms in *BMP15* and *GDF9* genes and investigating their association with litter size in Bangladeshi goat. The findings of the study reveal that different genotypes at three loci in *BMP15* gene had significant (p ≤ 0.05) effect on litter size in the prolific Black Bengal breed of Bangladesh. Hence, there is a need for further research with a substantially large number of animals across a wide range of geographically divergent populations of this breed. Our results enrich the repository of molecular markers database of caprine fecundity genes which pave the way for association studies with fecundity trait, hence contribute to molecular breeding in goat.

## Acknowledgement

The study was supported financially by a grant from the University Grants Commission Bangladesh under a project entitled “Screening genetic markers in fecundity: a savings account for marker-assisted selection in Black Bengal goat’.

## Data availability

The data that support the findings of this study are available from the authors upon reasonable request.

## Author contributions

A. Das conceived this project. M. Shaha performed sample collection. M. Shaha and A. Dutta extracted DNA and performed PCR. A. Das and M. Das Gupta performed data curation and formal data analysis. A. Das and O. F. Miazi involved in funding acquisition, project administration and supervision. A. Das drafted the original manuscript. All authors read and approved the final manuscript.

## Statement of animal rights

The manuscript does not contain clinical studies or patient data.

## Competing Interests

The authors declare that they have no competing interests.

## Notes

### Competing Interest Statement

The authors have declared no competing interest.

## References

Abdoli, R., Mirhoseini, S.Z., Hossein-Zadeh, N.G. and Zamani, P., 2018. Screening for causative mutations of major prolificacy genes in Iranian fat-tailed sheep. International Journal of Fertility and Sterility, 12, 51–55.

Adzhubei, I., Jordan, D.M. and Sunyaev, S.R., 2013. Predicting functional effect of human missense mutations using PolyPhen-2. Current Protocols in Human Genetics, 07, unit, 7.20.

Ahlawat, S., Sharma, R. and Maitra, A., 2013. Screening of indigenous goats for prolificacy associated DNA markers of sheep, Gene, 517, 128–131.

Ahlawat, S., Sharma, R., Roy, M., Mandakmale, S., Prakash, V. and Tantia, M.S., 2016. Genotyping of Novel SNPs in BMPR1B, BMP15, and GDF9 Genes for Association with Prolificacy in Seven Indian Goat Breeds. Animal Biotechnology, 27, 199–207.

Ahlawat, S., Sharma, R., Roy, M., Tantia, M.S. and Prakash, V., 2015. Association analysis of novel SNPs in BMPR1B, BMP15 and GDF9 genes with reproductive traits in Black Bengal goats. Small Ruminant Research, 132, 92–98.

Altschul, S.F., Gish, W., Miller, W., Myers, E.W. and Lipman, D.J., 1990. Basic local alignment search tool. Journal of Molecular Biology, 215, 403–410.

Amin, M., Husain, S. and Islam, A., 2000. Evaluation of Black Bengal goats and their cross with the Jamunapari breed for carcass characteristics. Small Ruminant Research, 38, 211–215.

Chen, N., Juric, I., Cosgrove, E.J., Bowman, R., Fitzpatrick, J.W., Schoech, S.J., Clark, A.G. and Coop, G., 2019. Allele frequency dynamics in a pedigreed natural population. Proceedings of the National Academy of Sciences of the United States of America, 116, 2158–2164.

Choudhury, M., Sarker, S., Islam, F., Ali, A., Bhuiyan, A., Ibrahim, M. and Okeyo, A., 2012. Morphometry and performance of Black Bengal goats at the rural community level in Bangladesh. Bangladesh Journal of Animal Science, 41, 83–89.

Chu, M.X., Liu, Z.H., Jiao, C.L., He, Y.Q., Fang, L., Ye, S.C., Chen, G.H. and Wang, J.Y., 2007. Mutations in BMPR-IB and BMP-15 genes are associated with litter size in Small Tailed Han sheep (Ovis aries). Journal of Animal Science, 85, 598–603.

Davis, G.H., 2005. Major genes affecting ovulation rate in sheep. Genetics Selection Evolution, 37, Suppl. S11–23.

Faruque, S., Chowdhury, S., Siddiquee, N. and Afroz, M., 2010. Performance and genetic parameters of economically important traits of Black Bengal goat. Journal of the Bangladesh Agricultural University, 8(1), 67–78.

Ghoreishi, H., Fathi-Yosefabad, S., Shayegh, J. and Barzegari, A., 2019. Identification of mutations in BMP15 and GDF9 genes associated with prolificacy of Markhoz goats. Archives Animal Breeding, 62, 565–570.

Gootwine, E., 2020. Invited Review: Opportunities for Genetic Improvement toward Higher Prolificacy in Sheep. Small Ruminant Research, 186:106090.

Gorlov, I.F., Kolosov, Y.A., Shirokova, N.V., Getmantseva, L.V., Slozhenkina, M.I., Mosolova, N.I., Bakoev, N.F., Leonova, M.A., Kolosov, A.Y. and Zlobina, E.Y.J.R.L.S.F.E.N., 2018. GDF9 gene polymorphism and its association with litter size in two Russian sheep breeds. Rendiconti Lincei. Scienze Fisiche e Naturali, 29, 61–66.

Hadizadeh, M., Mohammadbadi, M.R., Niazi, A., Esmailizadeh, A. and Gazooei, Y.M., 2014. Search for polymorphism in growth and differentiation factor 9 (GDF9) gene in prolific Beetal and Tali goats (Capra hircus). Journal of Biodiversity and Environmental Sciences, 4, 186–191.

Haldar, A., Pal, P., Datta, M., Paul, R., Pal, S.K., Majumdar, D., Biswas, C.K. and Pan, S., 2014. Prolificacy and its relationship with age, body weight, parity, previous litter size and body linear type traits in meat-type goats. Asian-Australasian Journal of Animal Sciences, 27(5), 628–634.

Hanrahan, J.P., Gregan, S.M., Mulsant, P., Mullen, M., Davis, G.H., Powell, R. and Galloway, S.M., 2004. Mutations in the genes for oocyte-derived growth factors GDF9 and BMP15 are associated with both increased ovulation rate and sterility in Cambridge and Belclare sheep (Ovis aries). Biology of Reproduction. 70, 900–909.

Heikal, H. and El Naby, W., 2017. Genetic improvement of litter size in four goat breeds in Egypt using polymorphism in bone morphogenetic protein 15 gene. Advances in Animal and Veterinary Sciences, 5, 410–415.

Husain, S., Amin, M., Islam, A., 1998. Goat production and its breeding strategy in Bangladesh. First National Workshop on Animal Breeding. Bangladesh Agricultural University, Mymensingh, pp. 17–36.

Jalbani, M.A., Kaleri, H.A., Baloch, A.H., Bangulzai, N., Bugti, A.G., Ashraf, F., Kaleri, R.R., Jan, M., Bugti, G.A. and Khosa, A.N., 2017. Study of BMP15 gene polymorphisim in Lehri goat breed of Balochistan. Journal of Applied Environmental and Biological Sciences, 7, 84–89.

Javanmard, A., Azadzadeh, N. and Esmailizadeh, A.K., 2011. Mutations in bone morphogenetic protein 15 and growth differentiation factor 9 genes are associated with increased litter size in fat-tailed sheep breeds. Veterinary Research Communications, 35, 157–167.

Kumar, S., Stecher, G., Li, M., Knyaz, C. and Tamura, K.J.M.b., 2018. MEGA X: molecular evolutionary genetics analysis across computing platforms. Molecular Biology and Evolution, 35, 1547–1549.

Lassoued, N., Benkhlil, Z., Woloszyn, F., Rejeb, A., Aouina, M., Rekik, M., Fabre, S. and Bedhiaf-Romdhani, S., 2017. FecXBara Novel BMP15 mutation responsible for prolificacy and female sterility in Tunisian Barbarine Sheep. BMC Genetics, 18, 43.

Maitra, A., Sharma, R., Ahlawat, S., Borana, K. and Tantia, M.S., 2016. Fecundity gene SNPs as informative markers for assessment of Indian goat genetic architecture. Indian Journal of Animal Research, 50, 349–356.

Mather, K., Jinks, J.L., 1977. Introduction to Biometrical Genetics, Chapman and Hall London.

Miah, G., Das, A., Bilkis, T., Momin, M.M., Uddin, M.A., Alim, M.A., Mahmud, M.S. and Miazi, O.F., 2016. Comparative study on productive and reproductive traits of Black Bengal and Jamnapari goats under semi-intensive condition. Scientific Research Journal, 4, 1–6.

Mishra, C., Rout, M., Mishra, S.P., Sahoo, S.S., Nayak, G. and Patra, R.C., 2017. Genetic polymorphism of prolific genes in goat-a brief review. Exploratory Animal and Medical Research, 7, 132–141.

Otsuka, F., McTavish, K.J. and Shimasaki, S., 2011. Integral role of GDF-9 and BMP-15 in ovarian function. Molecular Reproduction and Development, 78, 9–21.

Paul, R., Rahman, A., Debnath, S. and Khandoker, M., 2014. Evaluation of productive and reproductive performance of Black Bengal goat. Bangladesh Journal Animal Science, 43, 104–111.

Polley, S., De, S., Batabyal, S., Kaushik, R., Yadav, P., Arora, J.S., Chattopadhyay, S., Pan, S., Brahma, B., Datta, T.K. and Goswami, S.L., 2009. Polymorphism of fecundity genes (BMPR1B, BMP15 and GDF9) in the Indian prolific Black Bengal goat. Small Ruminant Research, 85, 122–129.

Shi, Y.Y. and He, L., 2005. SHEsis, a powerful software platform for analyses of linkage disequilibrium, haplotype construction, and genetic association at polymorphism loci. Cell Research, 15(2), 97–98.

Shokrollahi, B. and Morammazi, S., 2018. Polymorphism of GDF 9 and BMPR 1B genes and their association with litter size in Markhoz goats. Reproduction in Domestic Animals, 53, 971–978.

Vacca, G.M., Dhaouadi, A., Rekik, M., Carcangiu, V., Pazzola, M. and Dettori, M.L., 2010. Prolificacy genotypes at BMPR 1B, BMP15 and GDF9 genes in North African sheep breeds. Small Ruminant Research, 88, 67–71.

Wang, X., Yang, Q., Wang, K., Zhang, S., Pan, C., Chen, H., Qu, L., Yan, H. and Lan, X., 2017. A novel 12-bp indel polymorphism within the GDF9 gene is significantly associated with litter size and growth traits in goats. Animal Genetics, 48, 735.

Wang, X., Yang, Q., Zhang, S., Zhang, X., Pan, C., Chen, H., Zhu, H. and Lan, X., 2019. Genetic Effects of Single Nucleotide Polymorphisms in the Goat GDF9 Gene on Prolificacy: True or False Positive? Animals, 9, 886.

Williams, S.C., 2016. News Feature: Genetic mutations you want. Proceedings of the National Academy of Sciences of the United States of America, 113, 2554–2557.

Yue, C., Bai, W.L., Zheng, Y.Y., Hui, T.Y., Sun, J.M., Guo, D., Guo, S.L. and Wang, Z.Y.J.A.b., 2019. Correlation analysis of candidate gene SNP for high yield in Liaoning cashmere goats with litter size and cashmere performance. Animal Biotechnology, 19, 1–8.

